# Oncogenes, tumor suppressor and differentiation genes represent the oldest human gene classes and evolve concurrently

**DOI:** 10.1101/493627

**Authors:** A. Makashov, S.V. Malov, A.P. Kozlov

## Abstract

Earlier we showed that human genome contains many evolutionarily young or novel genes with tumor-specific or tumor-predominant expression. We suggested to call them *TSEEN* genes, i.e. <u>T</u>umor <u>S</u>pecifically <u>E</u>xpressed, <u>E</u>volutionarily <u>N</u>ew genes. In this paper we performed a study of the evolutionary ages of different classes of human genes, using homology searches in genomes of different taxa in human lineage. We discovered that different classes of human genes have different evolutionary ages and confirmed the existence of *TSEEN* gene classes. On the other hand, we found that oncogenes, tumor-suppressor genes and differentiation genes are among the oldest gene classes in humans and their evolution occurs concurrently. These findings confirm predictions made by our hypothesis of the possible evolutionary role of hereditary tumors.

## Introduction

We are interested in the possible evolutionary role of tumors. In previous publications [1 – 6] we formulated the hypothesis of the possible evolutionary role of hereditary tumors, i.e. tumors that can be passed from parent to offspring. According to this hypothesis, hereditary tumors were the source of extra cell masses which could be used in the evolution of multicellular organisms for the expression of evolutionarily novel genes, for the origin of new differentiated cell types with novel functions and for building new structures which constitute evolutionary innovations and morphological novelties. Hereditary tumors could play an evolutionary role by providing conditions (space and resources) for the expression of genes newly evolving in the DNA of germ cells. As a result of expression of novel genes, tumor cells acquired new functions and differentiated in new directions, which might lead to the origin of new cell types, tissues and organs. The new cell type was inherited in progeny generations due to genetic and epigenetic mechanisms similar to those for pre-existing cell types [5].

Our hypothesis makes several nontrivial predictions. One of predictions is that tumors could be selected for new functional roles beneficial to the organism. This prediction was addressed in a special work [5, 7], in which it was shown that the “hoods” of some varieties of gold fishes such as Lionhead, Oranda, etc. are benign tumors. These tumors have been selected by breeders for hundreds of years and eventually formed a new organ, the “hood”.

The other prediction of the hypothesis is that evolutionarily young and novel genes should be specifically expressed in tumors. This prediction was verified in a number of papers from our laboratory [6, 8 – 17]. We have described several evolutionarily young or novel genes with tumor-predominant or tumor-specific expression, and even the evolutionary novelty of the class of genes – cancer/testis genes – which consists of evolutionary young and novel genes expressed predominantly in tumors (reviewered in [6]). We suggested to call such genes *TSEEN* genes, i.e. <u>T</u>umor <u>S</u>pecifically <u>E</u>xpressed, <u>E</u>volutionarily <u>N</u>ew genes [5; 6].

In this paper, we performed a systematic study of the evolutionary ages of different functional classes of human genes in order to verify one more nontrivial prediction of the hypothesis of the possible evolutionary role of hereditary tumors, i.e. the prediction of concurrent evolution of oncogenes, tumor suppressor genes and differentiation genes [2; 3; 5].

## Results

The curves of gene age distribution for different classes of human genes obtained by the ProteinHistorian tool are represented in Figures 1-7.

**Figure 1.**
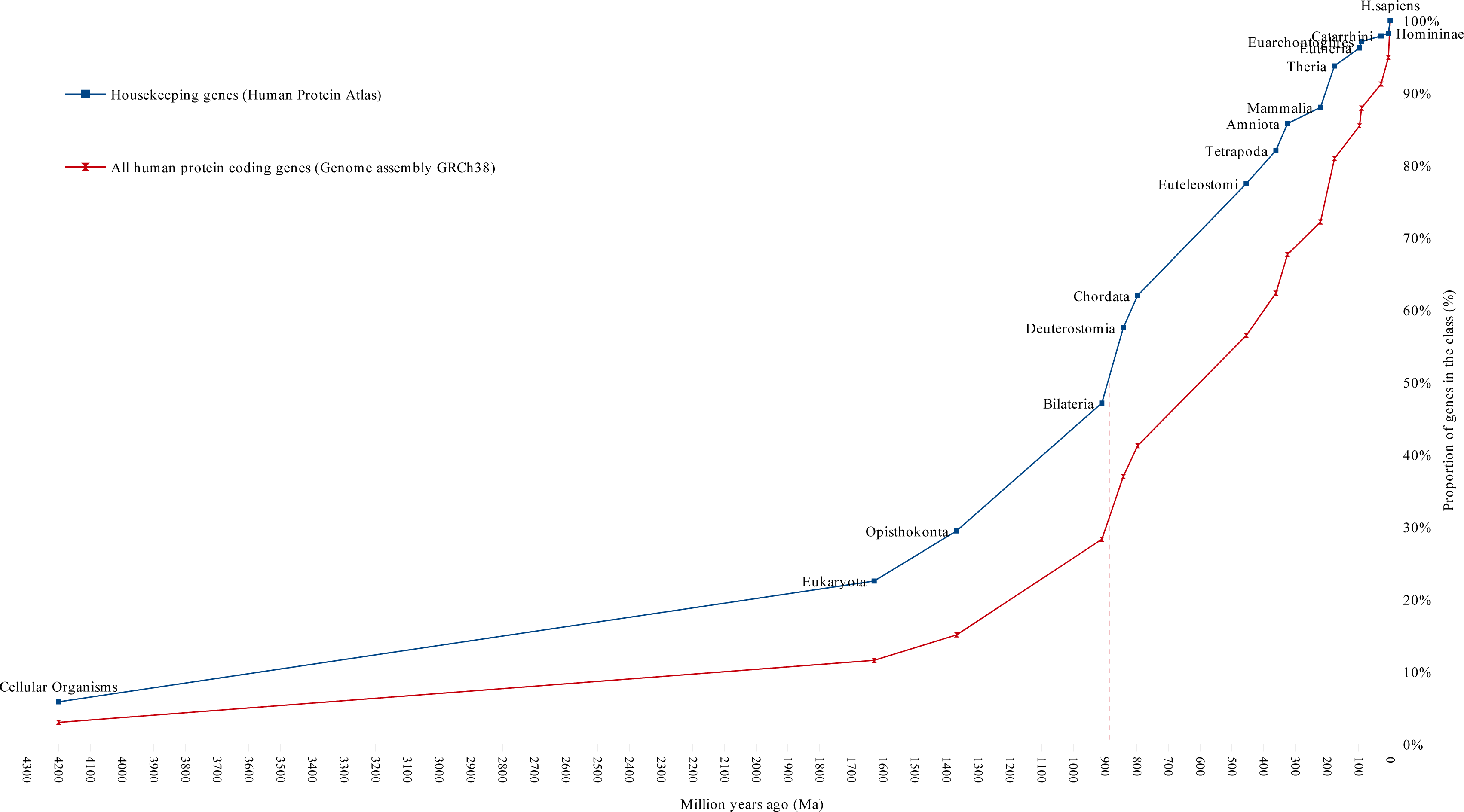
Distribution of human housekeeping genes genes and all protein coding genes according to their evolutionary ages. The evolutionary ages of the gene classes are measured numerically in million years at the median of distribution, i.e. at the time point on the human evolutionary timeline that corresponds to the origin of 50% of genes in this class.

These figures show curves sloping upward from left to right. The uppermost curve describes the gene age distribution of human housekeeping genes. The evolutionary age of this gene class, defined by the median position of the curve, is 894 Ma (Figure 1). The curve of all human protein-coding genes has evolutionary age of 600 Ma (Figures 1 and 3 - 5). This curve was used as control curve in our study. Some curves are located mainly above the control curve (Figure 3), others are located below it (Figures 4 and 5). The median ages of other groups of genes are the following: oncogenes (750 Ma), tumor suppressor genes (750 Ma), differentiation genes (693 Ma), homeobox genes (450 Ma), apoptosis genes (360 Ma), CT antigen genes (324 Ma), Biomedical Center globally subtracted, tumor-specifically expressed (BMC GSTSE) protein-coding genes (220 Ma), BMC GSTSE non-coding sequences (130 Ma), CT antigen genes located on X chromosome (CT-X) (60 Ma) and BMC GSTSE non-coding sequences located on X chromosome (BMC GSTSE-X non-coding sequences) (50 Ma) (Figure 2).

**Figure 2.**
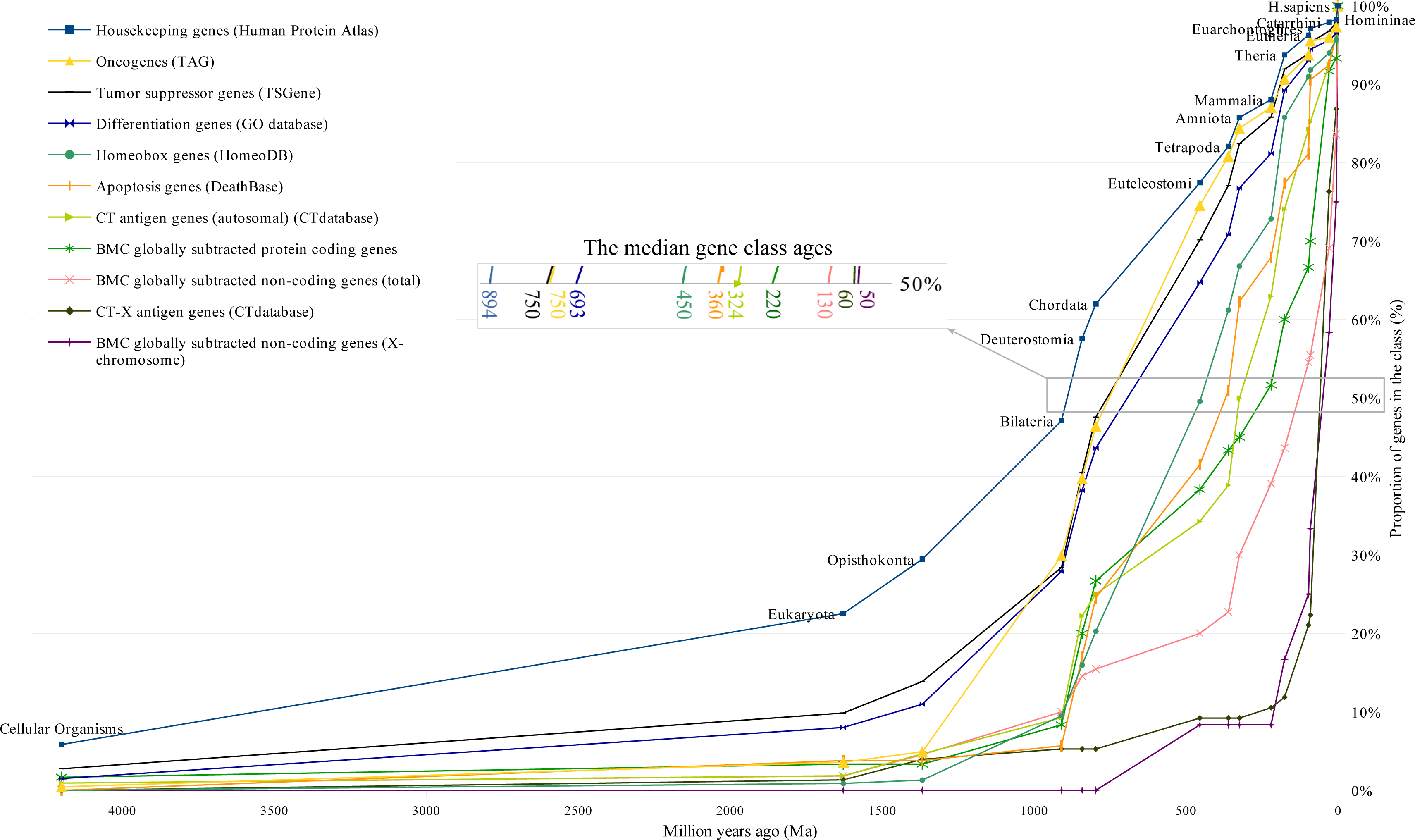
Gene age distributions of different classes of human genes

**Figure 3.**
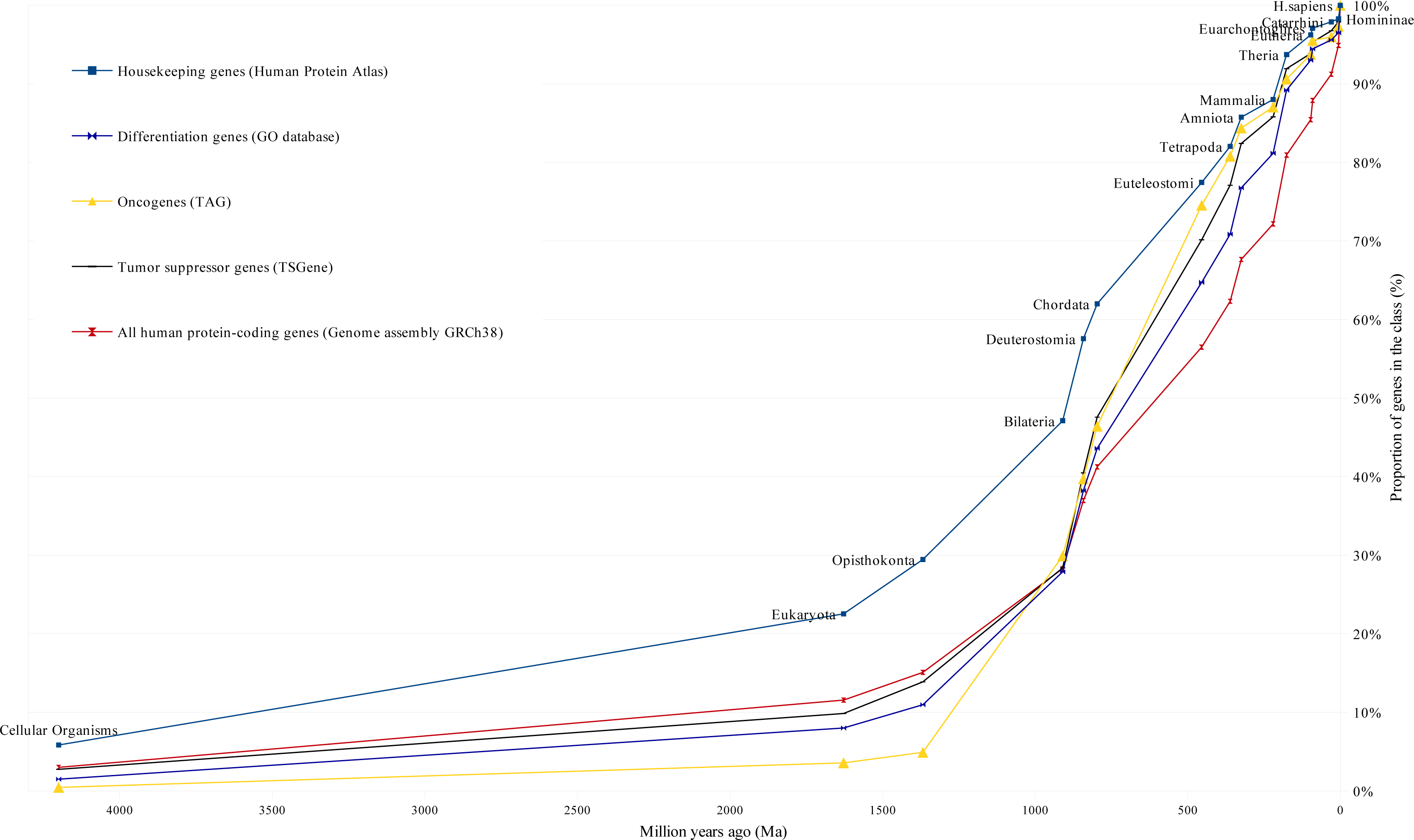
Cluster I of gene age distribution curves

**Figure 4.**
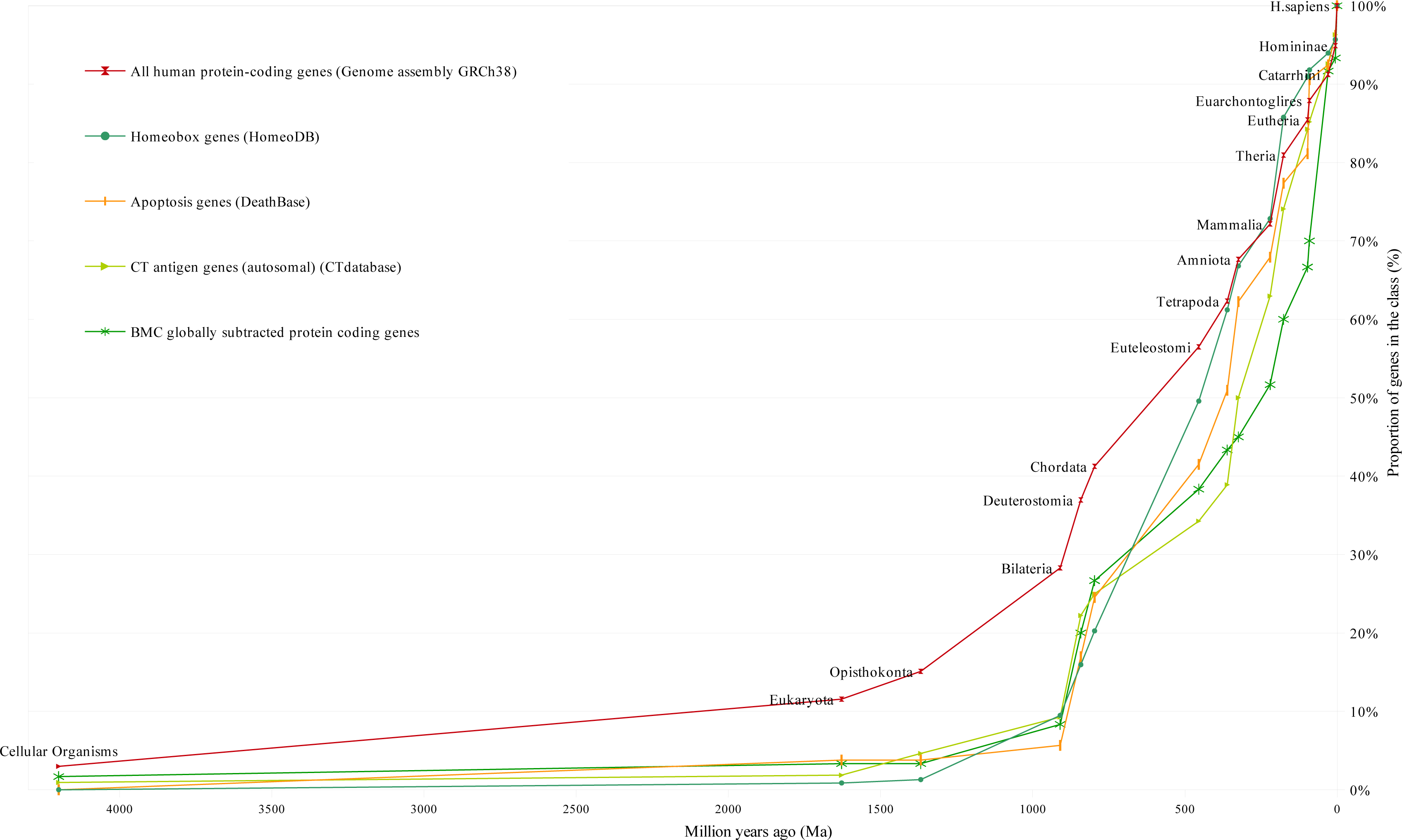
Cluster II of gene age distribution curves

**Figure 5.**
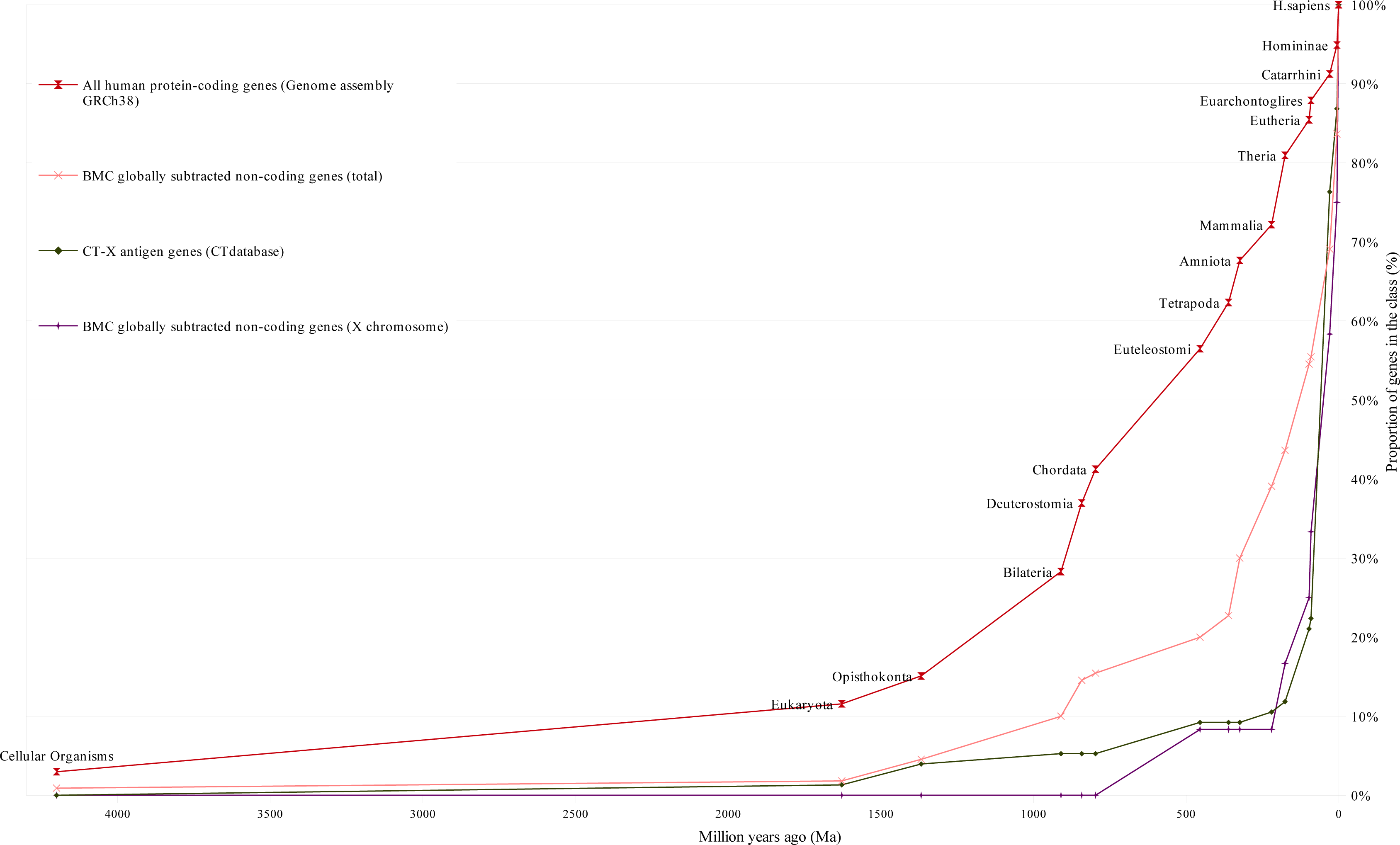
Cluster III of gene age distribution curves

As follows from Figures 2-5 the curves are organized in clusters. The existence of the clusters is supported by hierarchical cluster analysis (Figures 8 and 9, and Suppl. 2). The difference between the three clusters’ evolutionary ages is statistically significant (chi square P-value not exceed 1*10^-300^; X^2^= 1756 under 30 df) as well as the pairwise difference of the ages of each pair of clusters (Suppl. 2).

Cluster I includes the gene age distribution curves of human housekeeping genes, oncogenes, tumor suppressor genes and differentiation genes. It is located mainly above the control curve (Figure 3). Below the all protein-coding genes curve is the larger part of cluster II (including the following: homeobox genes, apoptosis genes, autosomal CT antigen genes and BMC GSTSE protein-coding sequences, Figure 4). The lowest position is occupied by cluster III, which includes curves of gene age distribution of the BMC GSTSE-X non-coding sequences and CT-X antigen genes orthologs (Figure 5).

The curves which belong to cluster I demonstrate growth starting from >4000 Ma. In *Bilateria* (910 Ma) they reach a proportion of 30%. The oncogene age distribution curve stays almost flat until *Opisthokonta* (1368 Ma), but after *Opisthokonta* goes upward and in *Bilateria* reaches 30% like other curves of cluster I. Between *Bilateria* and C*hordata* all curves of cluster I show a steep increase to 50%, and after *Chordata* (797 Ma) keep an almost constant slope up to 100% (Figures 2 and 3). The curve of housekeeping gene ages reaches 23% in *Eukaryota*, 29% in *Opisthokonta*, 47% in *Bilateria*, and makes a similar jump of 15% between *Bilateria* and *Chordata* (Figure1).

The curves of cluster II are slightly sloping until *Opisthokonta*, then slowly grow between *Opisthokonta* and *Bilateria*, and then demonstrate the 20% jump between *Bilateria* and *Chordata*, similarly to the curves of cluster I. The curve of homeobox genes ages, which belongs to cluster II, demonstrates almost constant slope between *Bilateria* and *Eutheria* (Figures 2, 4).

The curves of CT-X antigen genes and BMC GSTSE-X non-coding sequences are characterized by the highest growth (as compared to other curves) of 78% and 67%, respectively, during the last 90 mln years (Figures 2 and 5-6). The gene ages curve of BMC GSTSE-X non-coding sequences occupies the lowest position during the peri od of the last 67 Ma (53%) and and shows the maximum slope during the period of the last 6 Ma (25%), when the majority of other curves stop increasing (Figures 6 and 7).

**Figure 6.**
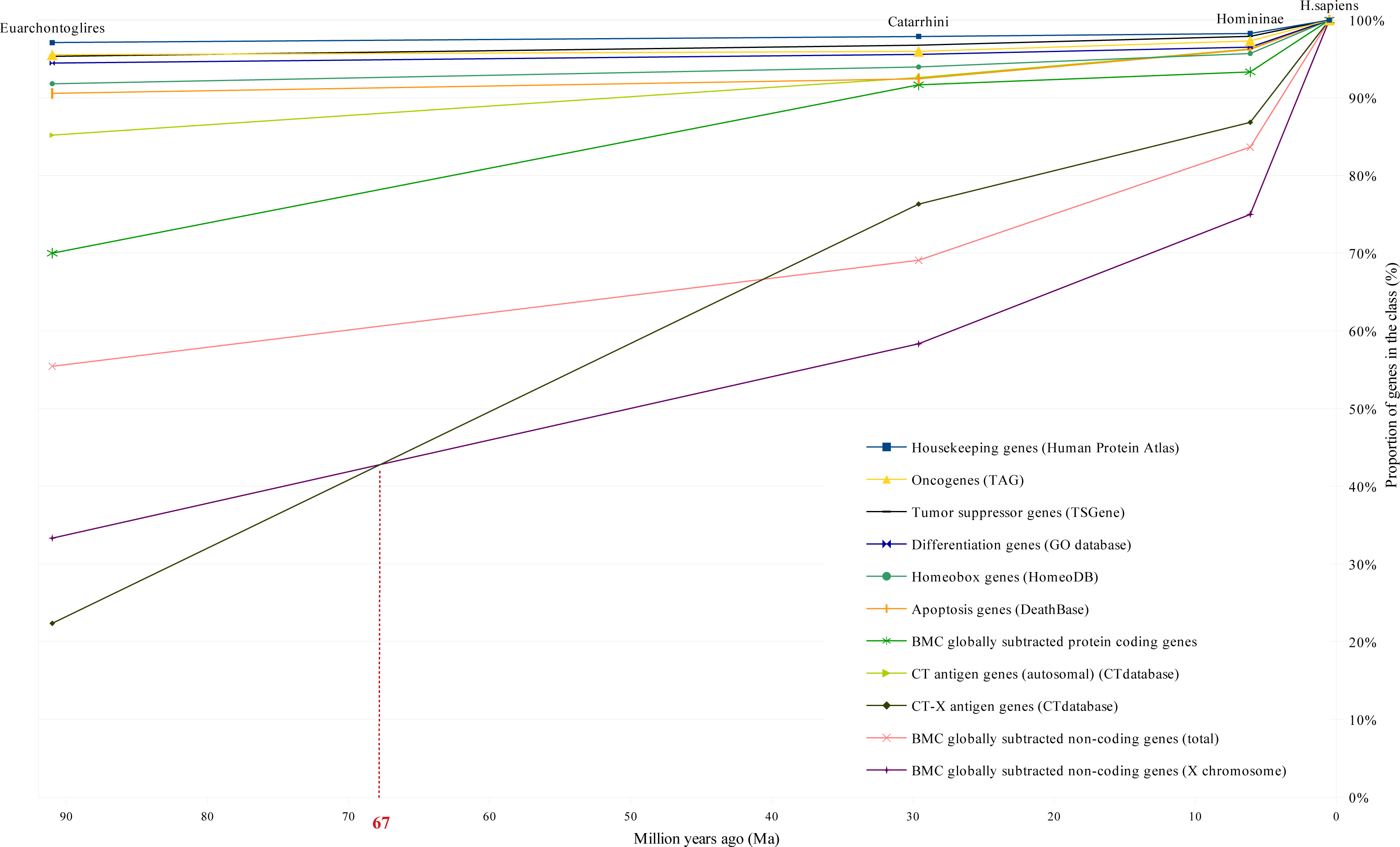
Gene age distribution for different classes of human genes between Euarchontoglires and H. sapiens

**Figure 7.**
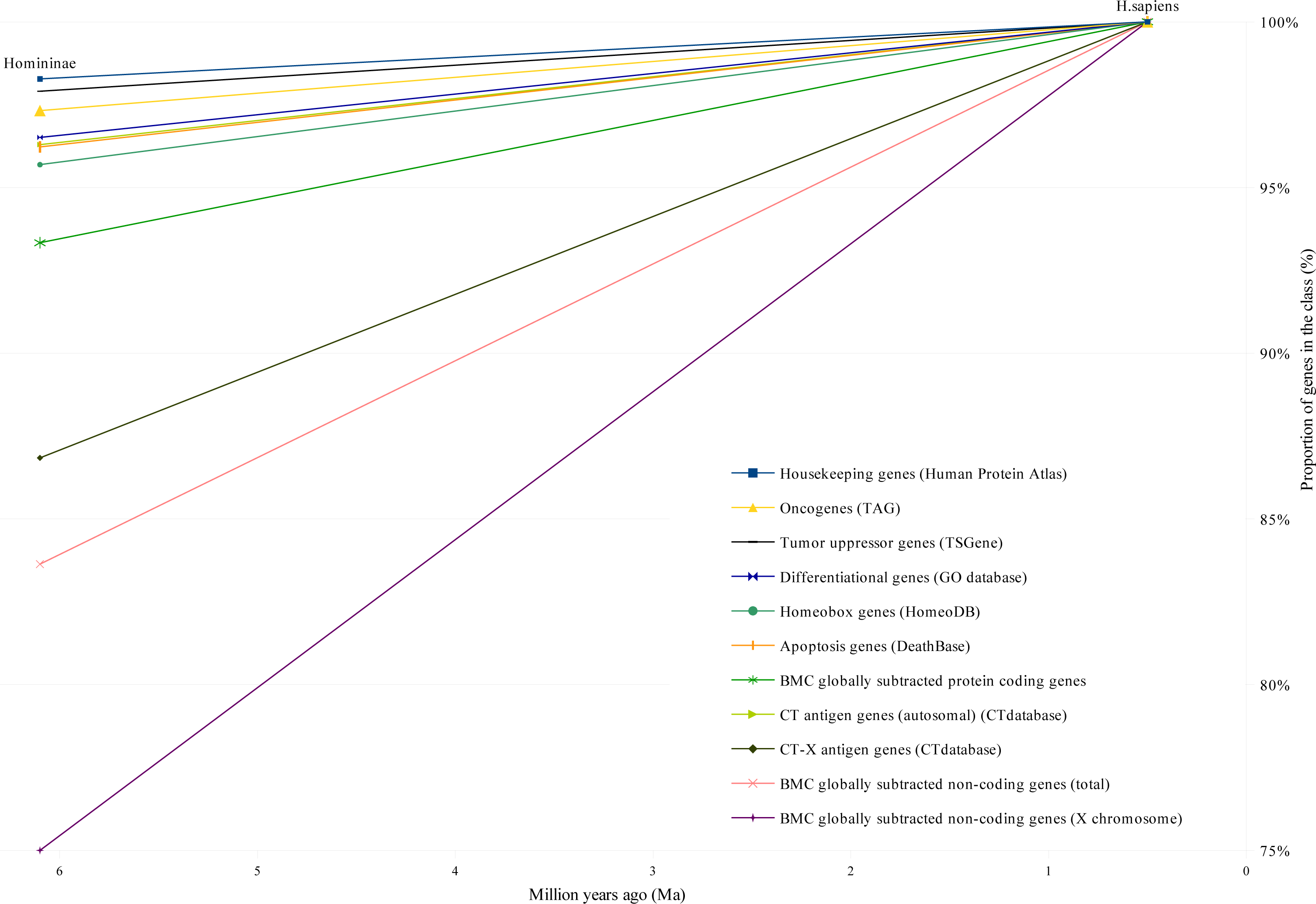
The proportion of different classes of human genes originated between Homininae and H. sapiens

**Figure 8.**
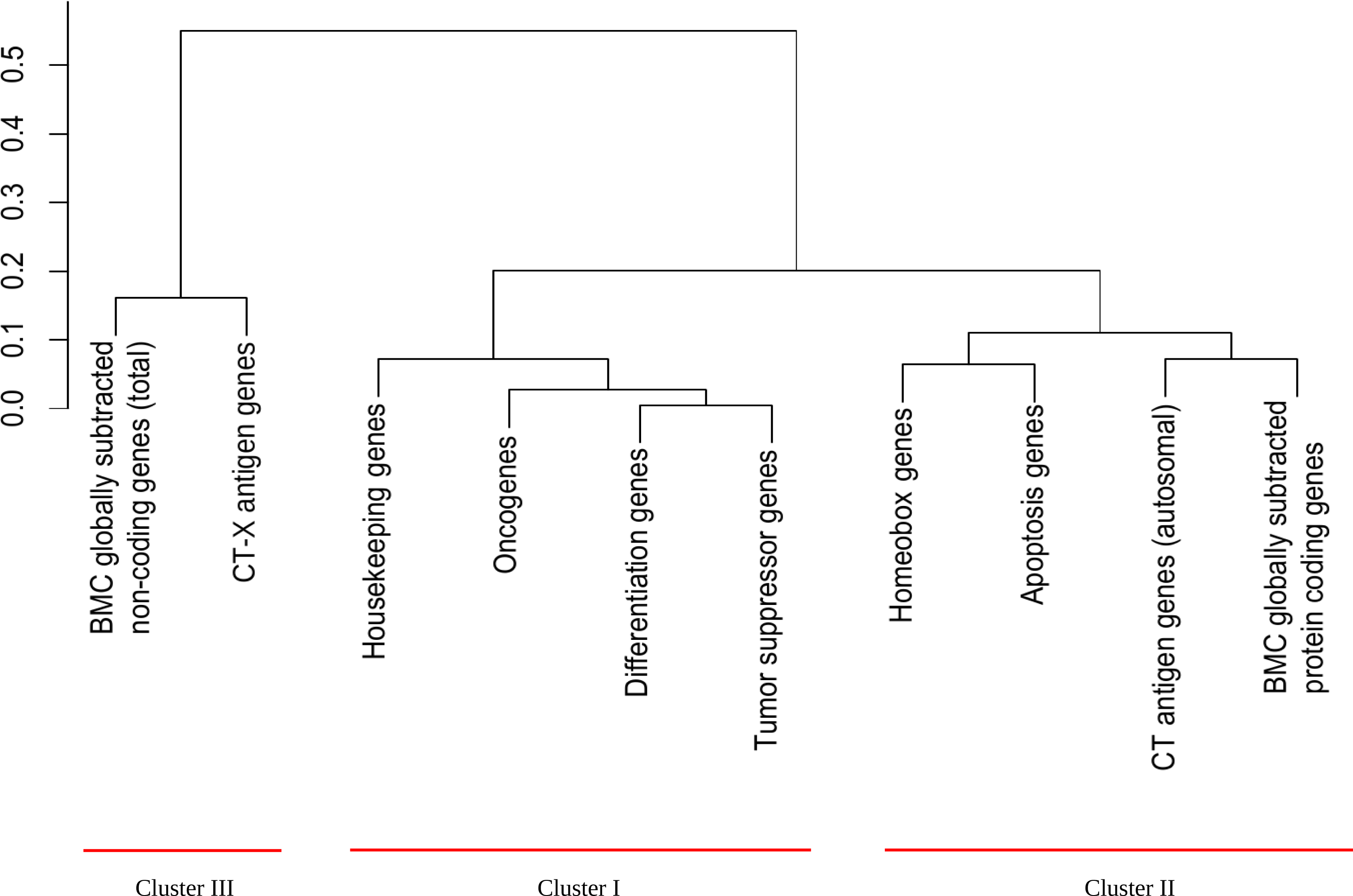
Hierarchical classification of different classes of human genes (Hellinger distance, complete linkage)

**Figure 9.**
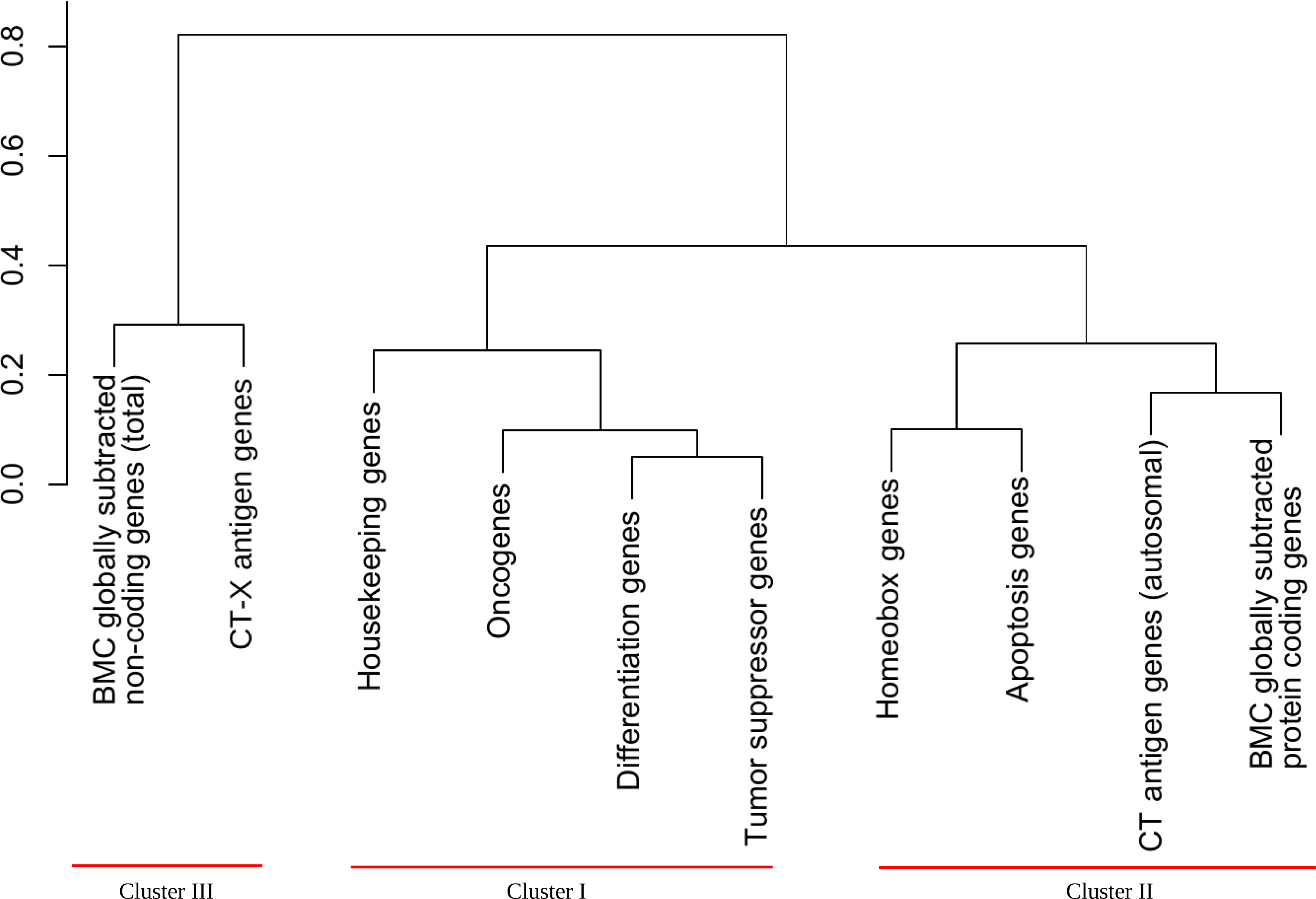
Hierarchical classification of different classes of human genes (Kolmogorov-Smirnov, complete linkage)

The CT-X antigen gene class was stochastically younger than the housekeeping gene class (two sided test P-value 0.027) and tumor suppressor gene class (two sided test P-value was 0.049), but after correction for multiple testing, simultaneously these results are not significant (see Suppl. 3 for complete pairwise relative evolutionary novelty analysis for different gene classes). Moreover, we discovered that the class of the BMC GSTSE non-coding sequences was stochastically younger than the class of housekeeping genes (two sided test P-value 2.6*10^-4^) and that the differentiation gene class was stochastically younger than the housekeeping gene class (two sided test P-value 5*10^-6^).

We also found that cluster III was stochastically younger than cluster I (two sided test P-value is 1.7*10^-5^) and the combination of clusters I and II (two sided test P-value is 1.9*10^-5^). Moreover, cluster III was stochastically younger than all protein-coding genes (P-value 0.0015) (Suppl. 4).

## Discussion

To study different functional classes of genes we used publicly available gene databases describing different gene classes – The Human Protein Atlas (housekeeping genes); TAG database (oncogenes); TSGene (tumor suppressor genes); CTDatabase (cancer/testis (CT) antigen genes); HomeoDB (HomeoBox genes); DeathBase (apoptosis genes); GeneOntology (differentiation genes); Biomedical Center Database (BMC GSTSE protein-coding genes and BMC GSTSE non-coding sequences). All annotated human protein coding genes (Genome assembly GRCh38) were used as control. Although we understand the limitations of such an approach connected with differing philosophies of the authors of databases and continuing upgrading of databases, we were able to obtain meaningful results. The results were also reproducible for different versions of databases with curves corresponding to different versions almost overlapping (see supplement 5).

We decided to study the ages of different gene classes in order to verify the predictions which stem from the hypothesis of the possible evolutionary role of heritable tumors formulated by one of us [5]. According to this hypothesis, hereditary tumors were the source of extra cell masses, which might be used in the evolution of multicellular organisms for the expression of evolutionarily novel genes and for the origin of new differentiated cell types with novel functions.

The evolutionary role of cellular oncogenes might consist in sustaining certain level of autonomous proliferative processes in the evolving populations of organisms and in promoting the expression of evolutionarily new genes. After the origin of a new cell type, the corresponding oncogene should have turned into a cell type-specific regulator of cell division and gene expression. If true, the number of cellular oncogenes should correspond to the number of cell types in higher animals [2; 3; 5].

If tumors and cellular oncogenes played a role in evolution as proposed, then the evolution of oncogenes, tumor suppressor genes, differentiation genes and cell types should proceed concurrently [5].

We found that any functional gene class includes genes with different evolutionary ages. This means that genes with similar functions originated during different periods of evolution. The age of a gene was defined by the most recent common ancestor on the human evolutionary timeline containing genes with similar sequences, i.e. with a significant BLAST score (or HMMER E-value).

The age of a functional gene class (or the age of the cluster) was described by distribution of ages of genes belonging to this gene class (i.e. particular gene database). For convenience, the age of the gene class can be measured numerically in million years at the median of distribution, i.e. at the time point on the human evolutionary timeline that corresponds to the origin of 50% of genes in this class. We found that different functional classes of human genes have different evolutionary ages ranging from 894 millions years for housekeeping genes to 50 million years for BMC GSTSE-X non-coding sequences. This reflects the different evolutionary history of different functional gene classes.

The curves of the older gene classes occupy the higher-left position and those of younger gene classes occupy the lower-right position on distribution curves (Figures 1-7). The slope of curves changes along the evolutionary timeline. This suggests that the rate of novel genes origin is different during different periods of evolution. Thus, the slope of all curves of clusters I and II, including the housekeeping gene ages distribution curve, increases sharply during the period between the origin of *Bilateria* and the origin of *Chordata* when many new cell types and morphological novelties originated. About 20% of all orthologs emerge during this period. Trends of the curves during the period of the Cambrian explosion (542 Ma), when most major animal phyla appeared in the fossil record [18], suggest that this radiation was preceded and followed by the extensive origin of novel genes (Figures 2-5). We see the last considerable increase in the origin of new genes 6 Ma ago, between *Homininae* and *H. sapiens*, when 15% of CT-X antigen genes, 10% of BMC GSTSE protein-coding genes, 17% of BMC GSTSE non-coding sequences and 25% of BMC GSTSE-X non-coding sequences originated (Figures 6 and 7).

It is known that housekeeping genes represent the oldest gene class in existing cells and evolve more slowly (according to their Ka/Ks rates) than tissue-specific genes [19; 20]. We found that the class of human housekeeping genes as described previously in [21] also contains evolutionarily younger genes, i.e. housekeeping genes continue to originate in the course of evolution, although at relatively slower rate than genes in other functional gene classes (see the slope of the corresponding curve). But as far as the class of housekeeping genes is large (7367 genes according to Uhlen et al. [21], even in humans 117 housekeeping genes originated, according to our data.

The intensive increase in the number of oncogenes began between *Opisthokonta* and *Bilateria* (25% of oncogenes), which coincided with the origin of multicellularity. This suggests a role for oncogenes in the origin of multicellular organisms. The other important jumps in the origin of oncogenes occur between *Bilateria* and *Chordata* (26%) and between *Chordata* and *Euteleostomi* (30%), which were periods of great morphological changes. Thus 83% of oncogenes originated between *Opisthoconta* and *Mammalia*.

Our data correspond with results of phylostratigraphic tracking of cancer genes which suggest a link to the emergence of multicellularity [22]. But our data also show considerable increase in the proportion of oncogenes and tumor suppressor genes before and beyond the emergence of vertebrates (Figures 2 and 3), while Domaset-Loso and Tautz described significantly lower origination of founder genes related to cancer beyond the emergence of vertebrates. This difference may be due to difference in methodology: Domaset-Loso and Tautz studied the emergence of cancer related domains while ProteinHistorian tool, which we used, studies the origin of the full-size proteins, in our case oncoproteins and tumor suppressor proteins.

While the origin of oncogene class, according to our data, is related to the origin of multicellularity, many differentiation genes were co-opted from unicellular ancestors (Figure 3). Today, genes that control metazoan development and differentiation are found in *Opisthokonta* suggesting that multicellularity evolved from unicellular opisthokont ancestors [23; 24; 25]. The slope of the differentiation gene ages distribution curve supports this notion. According to our data, 11% of human differentiation genes are conserved in *Opisthokonta* (Figure 3).

The gene classes studied in this paper form three clusters based on hierarchical cluster analysis. Each cluster contains curves with the least difference in gene age distributions.

The first cluster includes gene age distribution curves of housekeeping genes, oncogenes, tumor suppressor genes, and differentiation genes. This cluster is the oldest with evolutionary ages of gene classes from 894 Ma (housekeeping genes) to 693 Ma (differentiation genes). It is not homogeneous because the curve of housekeeping gene ages is separate from the other curves of the cluster, and differentiation gene class is stochastically younger than housekeeping gene class. On the other hand, gene age distribution curves of oncogenes, tumor suppressor genes and differentiation genes almost overlap.

It was known for a long time that there are oncogenes, which are very ancient [26 – 30]. But to our knowledge this paper is the first indication in the literature that oncogenes represent the most ancient class of genes in human genome with the exception of housekeeping genes. The other interesting piece of data is that tumor suppressor genes and differentiation genes coevolve with oncogenes. The fact that orthologs of oncogenes, tumor suppressor genes and differentiation genes belong to the same cluster and their distribution curves almost overlap means that they evolve concurrently, as predicted earlier [2; 3; 5]. This supports our hypothesis that hereditary tumors at early or intermediate stages of progression might participate in the evolutionary origin of new differentiated cell types [4; 5]. Our prediction that there should be a general correspondence between the number of oncogenes and the number of cell types is also supported by the other existing data. Thus, the TAG database, which we used in this study, currently contains 245 human oncogenes. Domaset-Loso and Tautz used other data sets (Sanger Cosmic, NCBI Entrez section in CancerGenes, the CancerGenes and the Network of Cancer Genes (NCG)). They found 380 oncogenes in these databases [22]. On the other hand, the current estimate of the number of the cell types in humans produced the number of 411 cell types, including 145 types of neurons [31]. That is, the general correspondence between the number of cell types and the number of oncogenes does exist, as was predicted in [2; 3]. It is noteworthy that when such correspondence was first predicted in 1987, only 20 oncogenes have been described [32], and by 1996 – only 70 oncogenes [33].

We further hypothesized that at least three different classes of genes are necessary for the origin of a new cell type in evolution: oncogenes, tumor suppressor genes, and evolutionarily novel genes, which determine a new function [5]. The existence of cluster I supports our hypothesis, although the number of protein coding tumor suppressor genes (TSGene database, 1018 genes) and differentiation genes (Gene Ontology, 3697 genes) is higher than the number of oncogenes (TAG database, 245 genes). The existence of cluster I also supports the differentiation theory of cancer [34]. According to this theory, cancer is abnormal programming of gene function during cell differentiation. The loss of tissue-specific functions (e.g. due to mutations of corresponding genes) is connected with tumors.

The second cluster occupies the intermediate position between cluster I and custer III with evolutionary ages of gene classes between 450 Ma (homeobox genes) and 220 Ma (BMC GSTSE protein-coding genes). Cluster II locates mainly below the control curve, i.e. the curve of all protein coding genes.

It is extremely interesting that in the evolutionary timeline the distribution curves of gene ages of homeobox and apoptosis genes are separated from those of differentiation genes by the period of several hundred millions years, i.e. evolutionarily the origin of genes responsible for differentiation and organogenesis are widely separated. Thus, before *Bilateria*, almost 30% of differentiation genes originated, and only 10% of homeobox genes. Half of differentiation genes originated at 643 million years, and half of homeobox genes – at 450 million years. In M*ammalia* 87% of differentiation genes and 73% of homeobox genes are represented. Indeed, the processes of differentiation and organogenesis are separated in evolution. For example, the thyriod gland was diffuse in the common ancestor of vertebrates and still has a diffuse nature and lacks the capsule in cyclostomes and in teleostean fishes [35 – 38]. In mammalians and humans diffuse endocrine system and diffuse, unencapsulated bundles of lymphatic cells still exist. Nevertheless, during certain periods of the evolutionary timeline the curves of cluster I and cluster II behave in a similar manner. E.g. between *Bilateria* and *Chordata* the curves of cluster I and cluster II demonstrate similar jump of about 20%, although in cluster II this jump starts from much lower level.

Finally, the third cluster is the youngest with evolutionary ages between 130 Ma and 50 Ma. This cluster includes gene classes expressed predominantly in tumors – CT-X genes and BMC GSTSE-X non-coding sequences. Genes belonging to this cluster continue to originate during last 90 Ma, and even during the last 6 Ma, as shown in Figures 6 and 7. The youngest during the last 6 Ma period are tumor-specifically expressed non-coding sequences located on X chromosome, discovered at the Biomedical Center by global subtraction of cDNAs of all known normal libraries from cDNAs of all known tumor libraries [8; 10].

We already described the evolutionary novelty of CT-X antigen gene class earlier [17]. Later other authors reproduced our results with appropriate reference to our original paper [39]. Here we confirmed the evolutionary novelty of CT-X gene class using the current upgraded database of CT genes – CTDatabase, and with another method – ProteinHistorian. In this paper, we also described the new class of *TSEEN* genes – BMC GSTSE ncRNA genes. In our other work we discovered a new long non-coding RNA (lncRNA) – *OTP-AS1* (*OTP*-antisense RNA 1) [40], which belongs to cancer/testis sequences.

Statistical analysis supported the existence of two classes of *TSEEN* genes – CT-X gene class and BMC GSTSE ncRNA gene class (Suppl. 1), and the existence of the whole class of *TSEEN* genes – cluster III was stochastically younger than the combination of two clusters I and II (Suppl. 4). The existence of cluster III was predicted by our hypothesis [5].

Thus at least three evolutionary categories of gene classes are expressed in human tumor cells: evolutionarily old (e.g. oncogenes), evolutionarily young or novel (e.g. CT-X genes and BMC GSTSE non-coding sequences) and intermediate age gene classes (e.g. BMC GSTSE protein-coding genes). But even evolutionarily older gene classes contain evolutionarily novel genes, for example, oncogenes *CT45A1* and *TBC1D3* [41 – 43] (see also discussion of evolutionarily novel housekeeping genes above). On the contrary, even evolutionarily younger gene classes contain evolutionarily older genes (10% of all genes in CT-X and BMC GSTSE-X ncRNA gene classes).

The data presented in this paper support and extend the concept of tumor-specifically expressed, evolutionarily novel *(TSEEN*) genes, formulated in [3 – 5], and confirmed in [6; 8 – 17]. From the data presented in this paper we can see that even different classes of genes (e.g. CT-X antigen genes and BMC GSTSE non-coding sequences) could be tumor-predominantly expressed and evolutionarily young or novel.

Thus the data presented in this paper confirm two predictions of our hypothesis of the possible evolutionary role of tumors, i.e. concurrent evolution of oncogenes, tumor suppressor genes and differentiation genes, and the existence of tumor specifically expressed, evolutionarily novel (*TSEEN*) gene classes. This may be important for better understanding of tumor biology, in particular of the possible evolutionary role of tumors as described in [5].

## Methods

The following public databases were used as a source of human gene classes in this study: housekeeping genes – The Human Protein Atlas; oncogenes – TAG database; tumor suppressor genes – TSGene; differentiation genes – GeneOntology; HomeoBox genes – HomeoDB; apoptosis genes – DeathBase; cancer-testis (CT) antigen genes – CTDatabase; BMC GSTSE protein-coding genes and non-coding sequences – Biomedical Center Database; and all annotated human protein coding genes – Genome assembly GRCh38 (21694 genes). CT antigen genes were divided into two groups: autosomal genes and genes located on X chromosome. BMC GSTSE non-coding sequences located on X chromosome were also separately studied.

Housekeeping genes are 7367 genes expressed in all analyzed tissues in the Human Protein Atlas [21]. This database contains information for a large majority of all human protein-coding genes regarding the expression and localization of the corresponding proteins based on both RNA and protein data. The Atlas contains information about 44 different human tissues and organs [21].

The TAG database (Tumor Associated Genes Database) (245 oncogenes) was designed to utilize information from well-characterized oncogenes and tumor suppressor genes to facilitate cancer research. All target genes were identified through text-mining approach from the PubMed database. A semi-automatic information retrieving engine collects specific information of these target genes from various resources and store in the TAG database. At the current stage, TAG database includes 245 oncogenes [44], which were used in our study. The database was modified for the last time on 2014.10.03.

TSGene 2.0 database contains 1217 human tumor suppressor genes (1018 coding and 199 non-coding genes) curated from a total of over 5700 PubMed abstracts [45]. In our study we used only 1018 protein-coding tumor suppressor genes.

Differentiation genes (3697 genes) were obtained by manual search for “differentiation” in the Gene Ontology database.

Homeobox gene database (HomeoDB) (333 genes) is a manually curated database of homeobox genes and their classification. HomeoDB2 includes all homeobox loci from 10 animal genomes (human, mouse, chicken, frog, zebrafish, amphioxus, nematode, fruitfly, beetle and honeybee) plus tools for downloading sequences, comparison between different species and BLAST search [46; 47]. We used the database, which was updated for the last time on 2011.08.08.

Deathbase (53 genes) is a database of proteins involved in different cell death processes. It is aimed to compile relevant data on the function, structure and evolution of this important cellular proccess in several organisms (human, mouse, zebrafish, fruitfly and worm). Information contained in the database is subject to manual curation [48]. The database was updated for the last time in 2011.

CTdatabase (276 genes) provides basic information including gene names and aliases, RefSeq accession numbers, genomic location, known splicing variants, gene duplications and additional family members. Gene expression at the mRNA level in normal and tumor tissues has been collated from publicly available data obtained by several different technologies. Manually curated data related to mRNA and protein expression, and antigen-specific immune responses in cancer patients are also available, together with links to PubMed for relevant CT antigen articles [49]. We used the update of 2017.

To construct the BMC database of sequences that are expressed in tumors but not in normal tissues, the normal EST set was subtracted *in silico* from the tumorous EST set. This approach is known as computer-assisted differential display (CDD). In total, 4564 cDNA libraries categorized as “tumorous” and 2304 “normal” libraries were used in CDD experiments. 251 EST clusters with tumor predominant expression were described in [8], and 196 clusters – in [10]. From these clusters 60 protein-coding genes and 121 non-coding sequences were selected for analysis.

All annotated human protein coding genes (21694 genes) were obtained from Genome assembly GRCh38 [50] with Ensembl tool [51]. The genome assembly was submitted on 2013.12.17.

The ProteinHistorian tool was used to perform homology search in genomes of different taxa.

The ProteinHistorian tool is an integrated web server, database and a set of command line tools which estimates the phylogenetic age of proteins based on a species tree, several external datasets of protein family predictions from the Princeton Protein Orthology Database (PPOD) [52] and two algorithms for ancestral family reconstruction (Dollo and Wagner parsimony) [53]. The ProteinHistorian tool searches the orthologs in 32 completely sequenced eukaryotic and prokariotic genomes from 16 taxa in the human ligeage (*Cellular Organisms, Eukaryota, Opisthokonta, Bilateria, Deuterostomia, Chordata, Euteleostomi, Tetrapoda, Amniota, Mammalia, Theria, Eutheria, Euarchontoglires, Catarrhini, Homininae,* and *H. sapiens*). The species tree used in analysis is presented in Supplement 6. Divergence time is estimated in millions of years ago (Ma) for each internal node in the species tree. It is important to note that a protein could have appeared at any time along the branch to which it is assigned, so the divergence time estimate reported is a lower bound. In addition, though the topology of this species tree is relatively non-controversial and the branch lengths reflect current research, the resulting age estimates are sensitive to the tree used. The ages are taken from the TimeTree database. Time tree database collects estimation of time of divergence among species data from publications in molecular evolution and phylogenetics. These included phylogenetic trees scaled to time (timetrees) and occasionally tables of time estimates and regular text. The data was collected from more than 2300 studies that have been published since 1987 [54].

The nucleotide BLAST algorithm, HMMER tool and the original Python script were used to analyze the ages of non-coding sequences. The orthologs were searched in 25 completely sequenced eukaryotic and prokaryotic genomes (Supplement 7).

The processing of datasets obtained with ProteinHistorian tool was carried out with Python script and Grep tool.

Some genes are included in several databases. This was taken into account in statistical analysis.

The age of the gene is defined by the most recent common ancestor on human evolutionary timeline containing genes with similar sequences, i.e. with a significant BLAST score (or HMMER E-value).

The age of the functional gene class (or cluster) is described by distribution of ages of genes belonging to this gene class. For convenience, the age of the gene class can be measured numerically in million years at the median of distribution, i.e. at the time point on the human evolutionary timeline which corresponds to the origin of 50% of orthologs of the functional gene class (Figure 1).

A probability distribution is stochastically smaller then another one if its cumulative distribution function is larger than the cumulative distribution function of the another one for each value of the argument. We say that a class of genes is stochastically younger than another one, if the age of this class is stochastically smaller than the age of the another class. Thus, we associate stochastically younger property of the gene class with its relative evolutionary novelty.

Before statistically analyze the relative evolutionarily novelty of gene classes we first evaluated stochastic difference in the age of gene classes using both Hellinger distance and Kolmogorov-Smirnov distance to specify clusters based on the complete linkage, and performed pairwise comparative statistical analysis by using the Kolmogorov-Smirnov and Chi-square tests to discover statistically significant differences between the evolutionary ages of gene classes.

We used appropriate contrasts and Sheffe S-method of multiple comparison to verify stochastic order in the evolutionarily ages of different genes classes observed in all the time points (taxons) from cellular organisms to humans. Thus, we apply covariance-adjusted method to create efficient joint confidence intervals for differences of the empirical distribution functions in all the time points available with the covariance obtained from the weak convergence of centered difference of the empirical distribution functions to the Brownian bridge process. The distribution of maximum modulus of correlated normal distributions required for the covariance-adjusted joint confidence interval was obtained by using Monte Carlo method with 10^6^ (before clustering) and 10^7^ (after clustering) replications.

## Supporting information

## Author Contribution statement

A.P.K. is an author of original hypothesis of the evolutionary role of tumors and *TSEEN* genes, and general design of experiments. A.M. performed bioinformatics analysis, and S.V.M. performed statistical analysis. All authors contributed to writing the manuscript.

## Acknowledgments

This study was supported by 5-100-2020 program grant at Peter the Great St. Petersburg Polytechnic University and a grant No833 from Ministry of Education and Science of the Russian Federation to A.P.K. We thank staff the Department of Information and Control Systems of Peter the Great St. Petersburg Polytechnic University for providing the access to processing power required to perform the analysis of the non-coding sequences. We thank D. Frishman, P. Drobintsev, and I. Selin for their help during this work.

## Abbreviations

BMC: Biomedical Center
BMC GSTSE: Biomedical Center globally subtracted, tumor-specifically expressed
BMC GSTSE-X non-coding sequences: Biomedical Center globally subtracted, tumor-specifically expressed non-coding sequences located on X chromosome
CDD: computer-assisted differential display
CT antigen gene: canser/testis antigen gene
*TSEEN* genes: tumor specifically expressed, evolutionarily novel genes
Ma: million years ago
PPOD: Princeton Protein Orthology Database
CT-X antigen genes: CT antigen genes located on X chromosome

