## Supplementary figures and images for "Oncogenes, tumor suppressor and differentiation genes represent the oldest human gene classes and evolve concurrently"

### Supplementary file 5

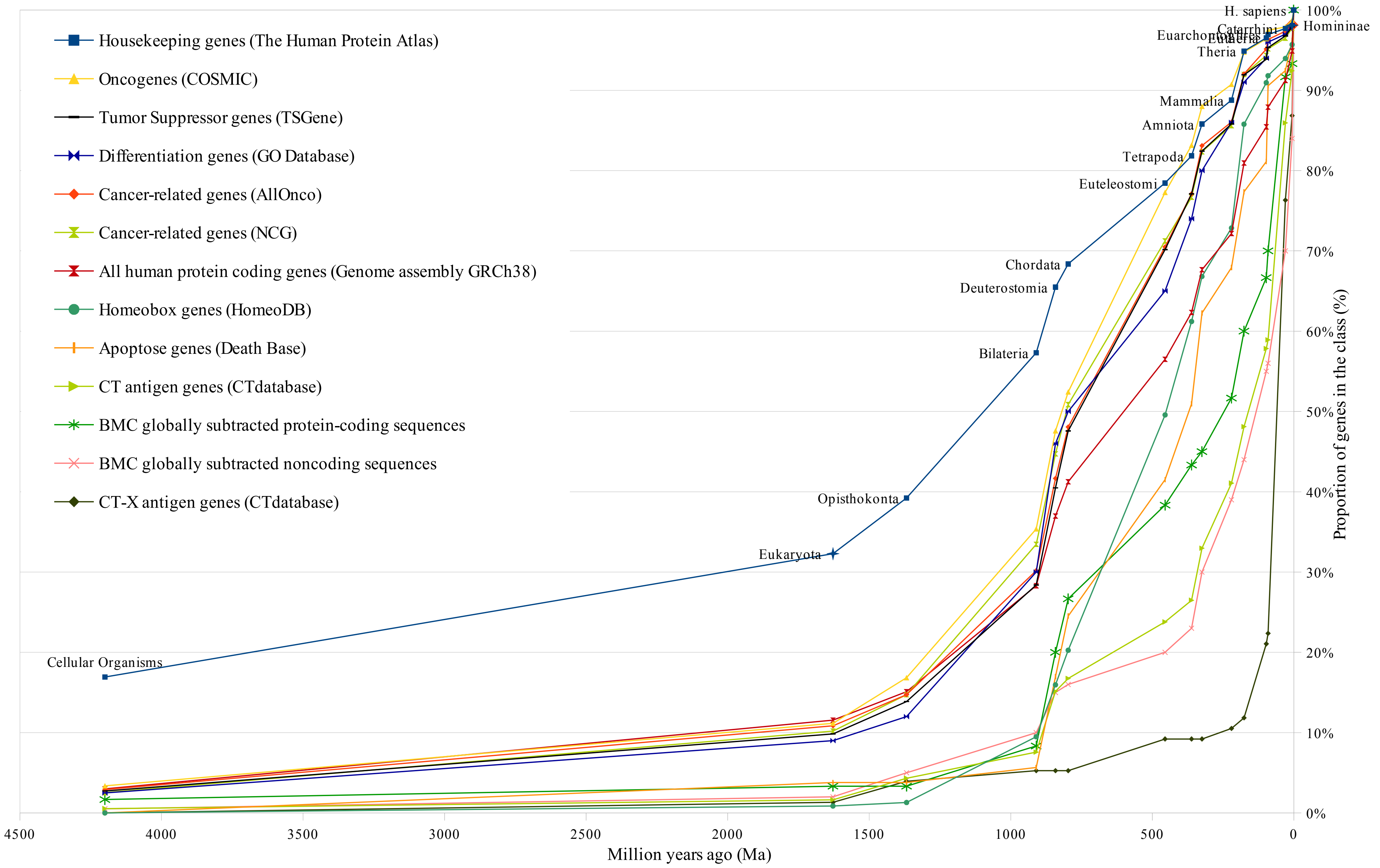


Supplement 5. One of previous versions of gene age distribution curves

### Supplementary file 6

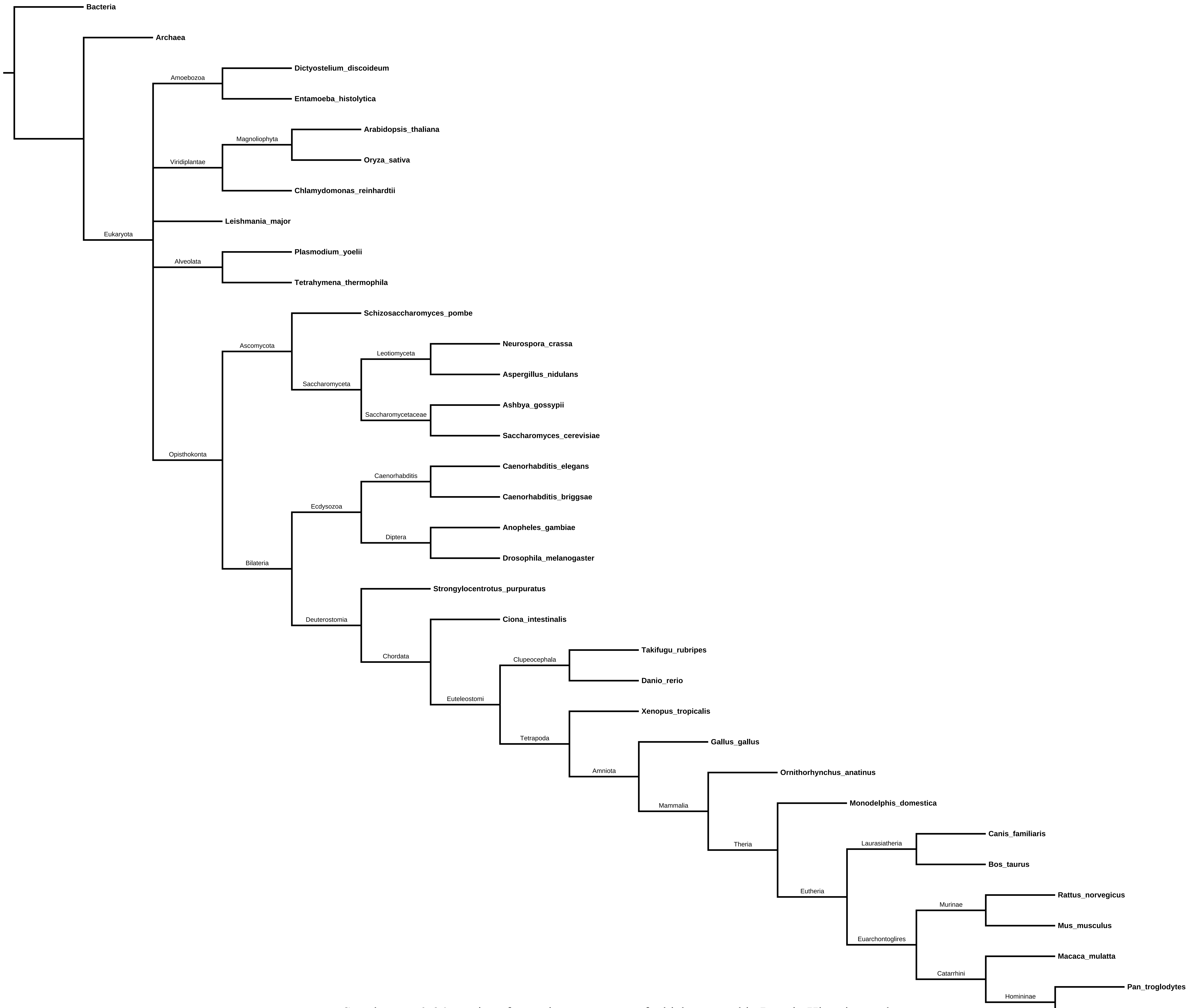


Supplement 6. 34 species of organisms genomes of which are used in Protein Historian tool
