## Supplementary material for "Oncogenes, tumor suppressor and differentiation genes represent the oldest human gene classes and evolve concurrently"

**Supplement 7. Genomes used for phylogenetic analysis of non-coding genes**

1. Escherichia coli str. K-12 substr. MG1655
2. Sulfolobus islandicus L.S.2.15
3. Plasmodium yoelii (assembly PY17X01)
4. Saccharomyces cerevisiae S288C (assembly R64)
5. Caenorhabditis elegans (assembly WBcel235)
6. Drosophila melanogaster (assembly Release 6 plus ISO1 MT)
7. Strongylocentrotus purpuratus (assembly Spur_4.2)
8. Ciona intestinalis (assembly KH)
9. Takifugu rubripes (assembly FUGU5)
10. Danio rerio (assembly GRCz11)
11. Xenopus tropicalis (assembly Xenopus_tropicalis_v9.1)
12. Crocodylus porosus (assembly CroPor_comp1)
13. Gallus gallus (assembly Gallus_gallus-5.0)
14. Ornithorhynchus anatinus (assembly Ornithorhynchus_anatinus-5.0.1)
15. Monodelphis domestica (assembly MonDom5)
16. Canis lupus familiaris (assembly CanFam3.1)
17. Bos taurus (assembly Bos_taurus_UMD_3.1.1)
18. Ovis aries (assembly Oar_v4.0)
19. Mus musculus (assembly GRCm38.p6)
20. Rattus norvegicus (assembly Rnor_6.0)
21. Macaca mulatta (assembly Mmul_8.0.1)
22. Nomascus leucogenys (assembly Nleu_3.0)
23. Pongo abelii (assembly Susie_PABv2)
24. Gorilla gorilla gorilla (assembly gorGor4)
25. Pan troglodytes (assembly Clint_PTRv2)
